# A Zika Virus Primary Isolate Induces Neuroinflammation, Compromises the Blood-Brain Barrier, and Upregulates CXCL12 in Adult Macaques

**DOI:** 10.1101/850198

**Authors:** Antonito T. Panganiban, Robert V. Blair, Julian B. Hattler, Diana G. Bohannon, Myrna C. Bonaldo, Blake Schouest, Nicholas J. Maness, Woong-Ki Kim

## Abstract

Zika virus (ZIKV) is a neurotropic virus that can cause neuropathy in adults and fetal neurologic malformation following infection of pregnant women. We used a nonhuman primate model, the Indian-origin Rhesus macaque (IRM), to gain insight into virus-associated hallmarks of ZIKV-induced adult neuropathy. We find that the virus causes prevalent acute and chronic neuroinflammation and chronic disruption of the blood-brain barrier (BBB) in adult animals. Infection results in significant, targeted, and sustained upregulation of the chemokine, CXCL12, in the central nervous system (CNS). CXCL12 plays a key role both in regulating lymphocyte trafficking through the BBB to the CNS, and in mediating repair of damaged neural tissue including remyelination. Understanding how CXCL12 expression is controlled will likely be of central importance in the definition of ZIKV-associated neuropathy in adults.

**Author summary:** Zika virus (ZIKV) is a virus that can cause neurological problems in adults and damage to the fetal brain. Nonhuman primates (NHPs) are usually superior animal models for recapitulating human neurological disease because their brain, nervous system structure and immune response to virus infection are very similar to that of humans. We have studied the effect of ZIKV infection on the adult NHP brain and made several significant observations. Infection resulted in a high incidence of mild to moderate brain inflammation that persisted for a surprisingly long period of time. We also found that the virus disrupted the blood brain barrier, which is important for controlling transport of material from blood to the brain. It appears that the central nervous system expresses a specific substance in response to virus infection called a chemokine. This specific chemokine may be involved in virus-induced inflammation and/or in repair of virus-induced brain damage. Our data are significant since they help in understanding the mechanism of brain damage caused by ZIKV in adults.

## Introduction

Zika virus (ZIKV), is a neurotropic flavivirus associated with Guillain-Barre’ syndrome (GBS) in adults and is also well-known for causing fetal neurologic malformation following infection of pregnant women (1, 2). In addition to causing GBS, which features damage to the protective myelin sheath surrounding axons, ZIKV can cause neuropathy in adults in the form of meningioencephalitis and myelitis (3, 4). The pathogenesis of ZIKV and the host-pathogen interactions important for the development of these lesions still need to be elucidated.

The blood-brain barrier (BBB), the boundary between circulatory and CNS tissues, is composed of brain microvascular endothelial cells (BMECs) and supporting associated pericytes and astrocytes. Intercellular tight junction (TJ) and adherens junction (AJ) integrity is important for maintenance of the intracellular network of MECs that comprises the vascular endothelium. Disruption of the BBB occurs during the pathogenesis of a wide range of infectious, autoimmune, and neurodegenerative diseases. Neurotropic flaviviruses, including Japanese Encephalitis virus (JEV) and West Nile virus (WNV), disturb the BBB in adults through disruption of the BMEC network (5, 6).

The chemokine CXCL12 is a key regulator of both myelin formation during embryogenesis and remyelination following neural damage in the CNS and peripheral nervous system (PNS) in adults (7, 8). CXCL12 facilitates the migration and maturation of oligodendrocyte precursor cells (OPCs) during CNS remyelination (7, 8). In the PNS, CXCL12 may similarly function in Schwann cell migration (9), as CXCL12 is expressed by perisynaptic Schwann cells during recovery of neuromuscular junctions following damage (10). In addition to serving as a key regulator of neural repair, CXCL12 plays an important function in regulating lymphocyte migration through the BBB. During homeostasis CXCL12 is expressed by BMECs resulting in spatial restriction of lymphocytes to the microvascular perivascular space, and changes in chemokine expression and distribution can lead to neuroinflammation(11, 12). The role of CXCL12 in neural repair, lymphocyte migration into the CNS parenchyma, and in multifarious functions during development and immunity, are mediated through interaction of this chemokine with its primary receptor, CXCR4 (13). Stimulation of CXCR4 by CXCL12 results in activation of an interwoven set of downstream effector pathways (14, 15).

Nonhuman primates (NHPs) are good animal models for recapitulating human neurological disease since they are genetically and physiologically similar to humans and exhibit CNS and PNS elaboration and brain morphology closely resembling that of humans. We have delineated the effect of ZIKV on the neural tissue of eighteen adult Indian Rhesus macaques (IRMs). Our data indicate that the virus causes acute and chronic inflammation of neural tissue in adult animals with accompanying damage to the BBB. Moreover, we find that expression of CXCL12 in the CNS is upregulated both during acute ZIKV infection and, surprisingly, for an extended time after infection. We propose that this chemokine is likely to be important in both long-term neuropathology and neural repair following ZIKV-induced damage of the PNS and CNS. Thus, this NHP model is valuable for experimentally deciphering important hallmarks of novel ZIKV-induced neuropathology in adults and in elucidating molecular mechanisms underlying virus-induced GBS spectrum disorders in humans.

## Results

### ZIKV infection causes acute and chronic perivascular neuroinflammation

We inoculated eighteen adult IRMs subcutaneously with 10^4^ plaque forming units (pfu)/animal of a minimally passaged Brazilian ZIKV isolate (Rio-U1)(16). An outline of the experimental design is shown in Fig. 1. Fifteen of the animals used in the study were females infected during pregnancy and three were adult males. The females were from a previous project focusing on the effect of ZIKV on the fetus (17). Following infection at different times during gestation, these animals all displayed acute viremia (Fig. S1), with some animals exhibiting transmission of virus to amniotic fluid. Infants were delivered by C-section, or pregnancy was experimentally terminated, or pregnancy ended through ZIKV-mediated demise. As outlined in Fig. 1, necropsy and collection of neural tissue was obtained from the dams at various times before or after parturition or fetal termination. Four of the dams (EL21, ID92, JR20, and JI20) were infected and necropsied 16 or 17 days after infection (acute), and the remainder were maintained for significantly longer periods of time after infection (3.5 to 10 months). The three adult males (HP17, HP87, and JP58) were infected and necropsied after 30 days.

**Figure 1.**
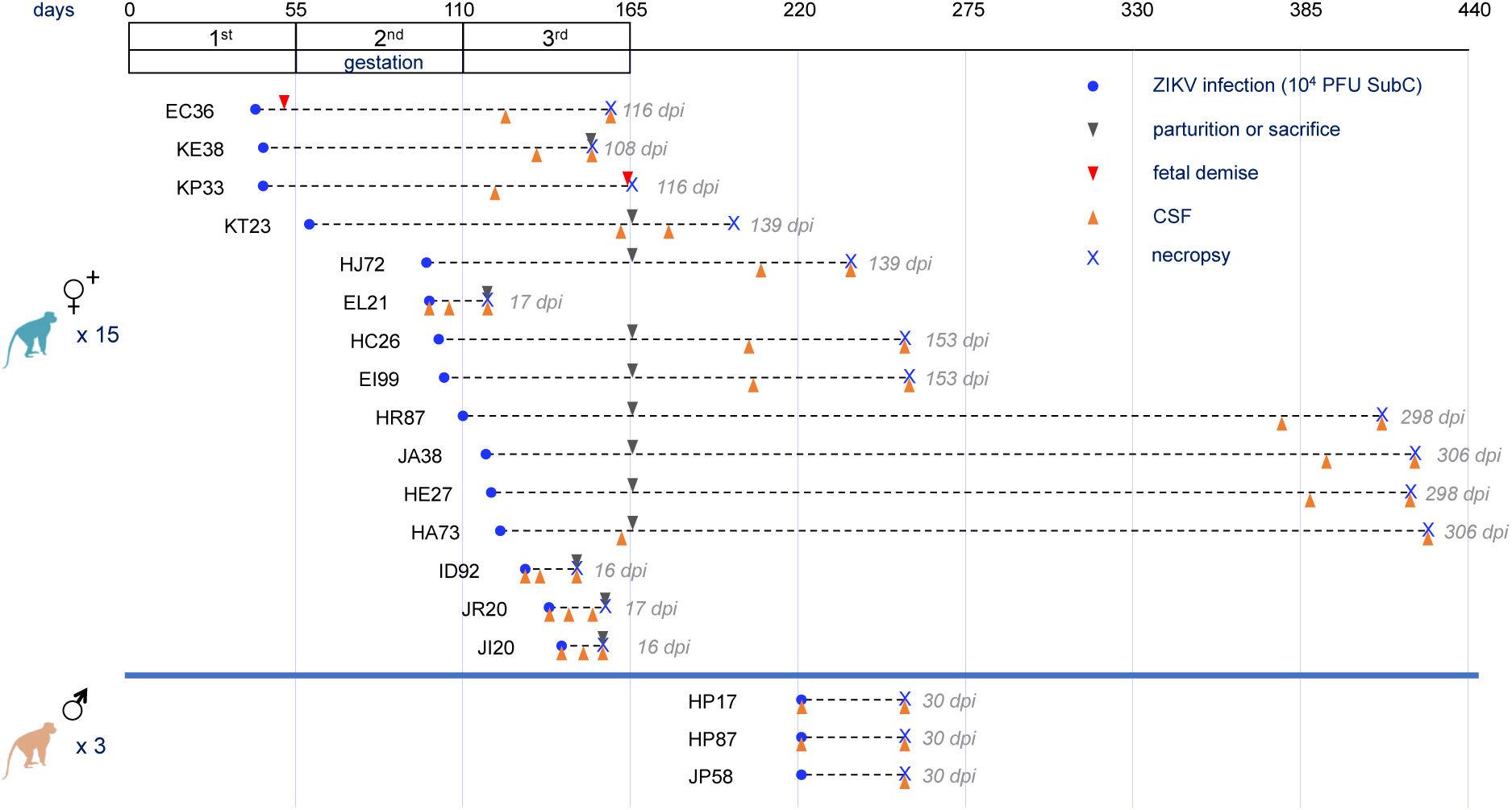
NHP study design. Fifteen female IRMs (17), and three male IRMs were infected with 10^4^ PFU of ZIKV strain Rio-1. Pregnancy in the Rhesus macaque is divided into three 55-day trimesters, which are equivalent in developmental landmarks to the trimesters in human pregnancy (55). These trimesters are depicted in the timeline at top, as are relative times of infection, sample collection, and necropsy for the dams. Acute viremia was detected in all animals (Figure S1).

Interestingly, H & E staining of CNS and PNS samples collected at necropsy revealed consistent scant to moderate perivascular inflammation in the meninges along with sporadic additional pathological features in the CNS parenchyma including glial nodules and lymphocytic infiltration (Fig. 2). Inflammation was observed in neural tissue both acutely and chronically after infection. Table 1 provides a summary of observations from neural tissue in individual animals.

**Table 1.** Summary of observed perivascular inflammation in the CNS and PNS of adult animals infected with ZIKV.

| Animal | Sciatic n. | Lumbar<br>cord | Thoracic<br>cord | Cervical<br>cord | Choroid<br>plexus | Brainstem | Cerebellum | Occipital<br>lobe | Temporal<br>lobe | Frontal<br>lobe | Basal Ganglia |
| --- | --- | --- | --- | --- | --- | --- | --- | --- | --- | --- | --- |
| EC36 |  |  | + |  |  |  |  |  |  |  |  |
| EL21 | + |  | + |  | + |  |  |  |  | + |  |
| HA73 | + |  |  |  |  |  |  |  |  |  |  |
| HE27 |  | + |  |  |  | ++ | ++ |  | + | ++ |  |
| ID92 |  | + | + | + |  |  |  |  |  |  |  |
| JA38 | + |  | + |  |  | + |  |  |  |  |  |
| JI20 | + |  |  |  |  |  |  |  |  |  |  |
| JR20 |  | + | + |  |  |  | + |  | + | + | + |
| KE38 |  |  | + |  |  |  |  |  |  |  |  |
| KT23 |  |  |  |  |  | + |  | + |  |  |  |
| JP58 |  | + |  |  |  |  |  |  |  |  |  |

**Figure 2.**
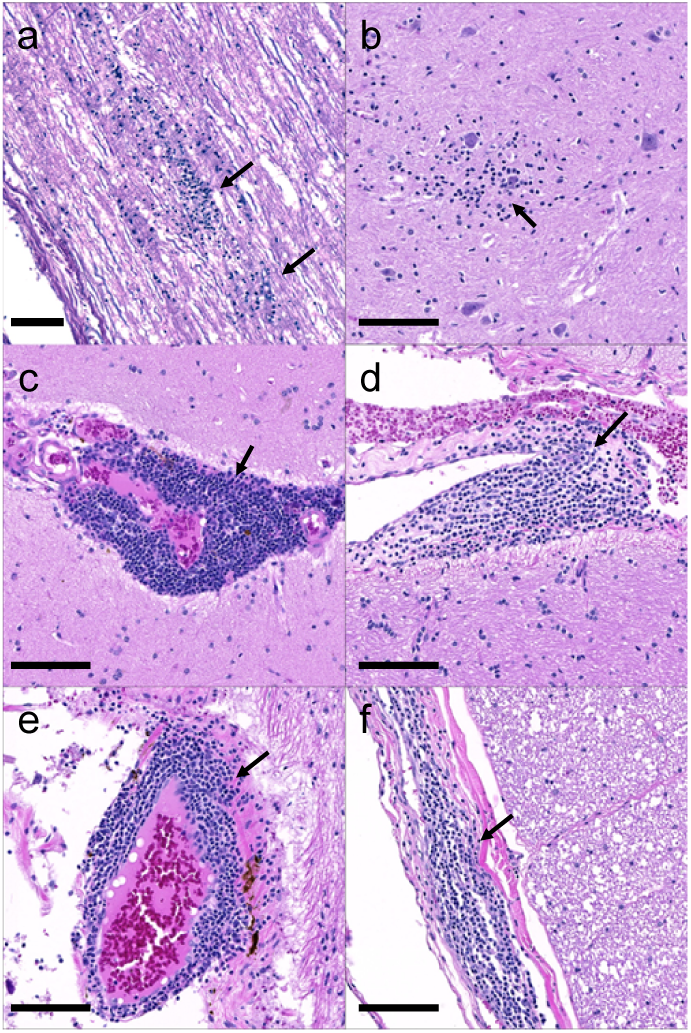
Neuroinflammation in the spinal cord of ID92 (A&B) and the spinal cord and brain of HE27 (C-F). ID92 had multifocal glial nodules (arrows) within both the white (a) and grey (b) matter of the spinal cord. HE27 had widespread perivascular inflammation (arrows) at multiple levels of the cerebrum (c&d), cerebellum (e), and spinal cord (f). H&E, Bar = 100um.

### ZIKV infection disrupts the adult blood-brain barrier

Since ZIKV caused perivascular inflammation with accompanying lymphocytic infiltration, this suggested that inflammation is likely to arise through BBB dysregulation. Fibrinogen is a protein normally restricted to serum. Thus, extravasation of fibrinogen into the perivascular space of CNS vessels is indicative of disruption of the BBB. To detect and quantify ZIKV-associated extravasated fibrinogen we performed multi-label immunofluorescence staining using anti-fibrinogen Ab. To visualize microvascular endothelial cells we used anti-GLUT-1 Ab. GLUT-1 is a plasma membrane protein found in abundance on vessel endothelial cells at the BBB (18) (Fig. 3a). Using dual overlay histograms of twenty five vessels per animal we quantified the percent of vessels with fibrinogen extravasation in ZIKV-infected and control groups in both cortical brain tissue and spinal cord (Fig. 3b). There was a significant increase in extravasated fibrinogen with infection in both brain and spinal cord (Fig 3c,d). This indicates that the integrity of the microvascular endothelium and the BBB was compromised following ZIKV infection.

**Figure 3.**
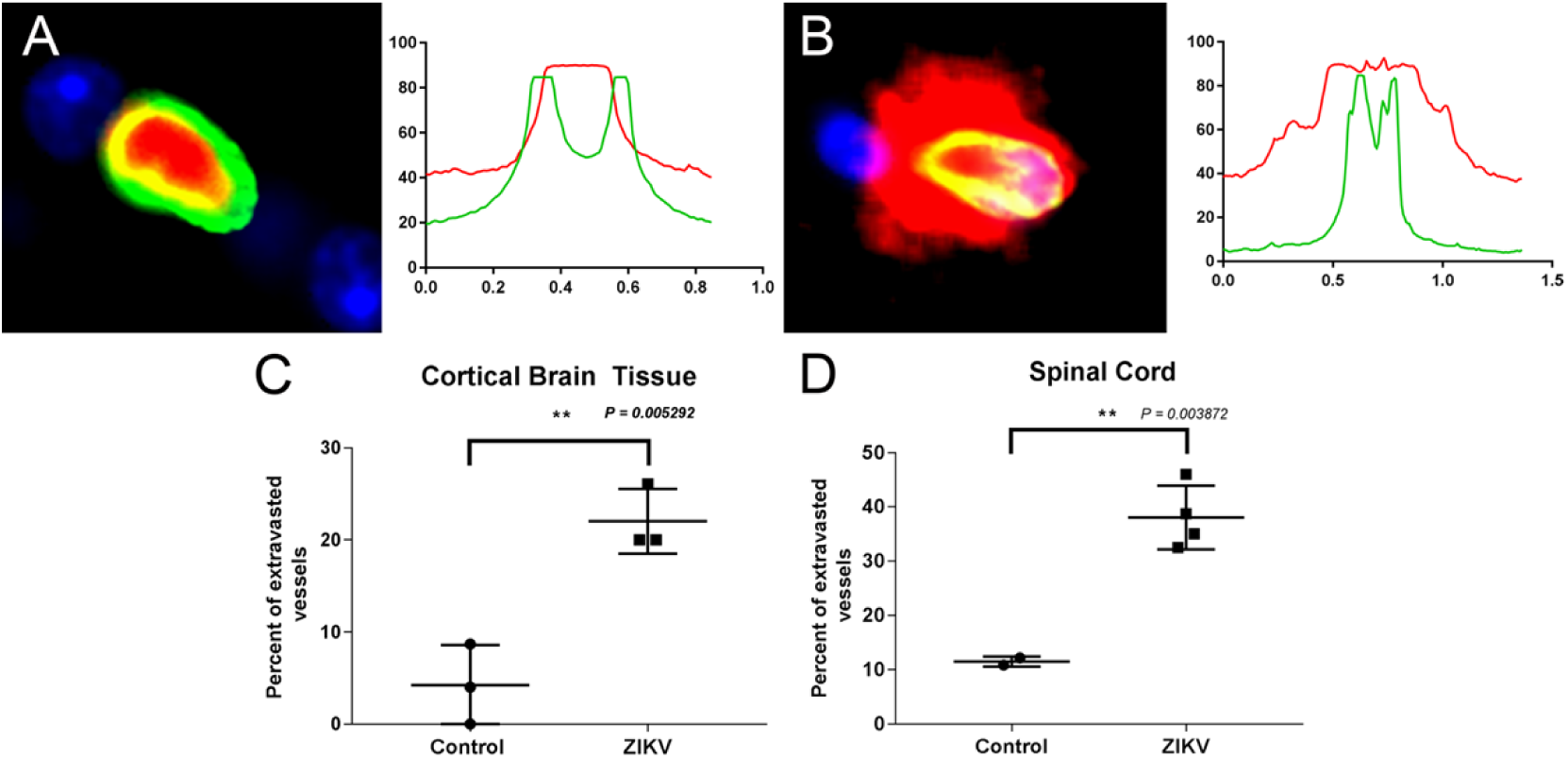
ZIKV disrupts the BBB in adult IRMs as evidenced by an increase in extravasated vessel fibrinogen during infection. Vessel endothelial cells were detected using Alexa Flour 488-labeled anti-GLUT-1 Ab (green) and fibrinogen detected using Alexa Flour 594-labeled anti-fibrinogen Ab (red). Nuclei were detected with DAPI. (A) Cross section and immunofluorescence analysis of a stained vessel without extravasation and associated dual overlay histogram. (B) Immunofluorescence stained vessel with extravasation and associated dual overlay histogram. Percent of extravasated vessels in (C) cortical brain tissue, (D and E) spinal cord.

### ZIKV infection acutely and chronically upregulates CXCL12 in the CNS

We then carried out a series of experiments focusing on components of the cerebral spinal fluid (CSF) from the infected animals to gain insight into the cause of ZIKV-induced neural inflammation in adult animals. In our prior study (17), efforts to detect ZIKV RNA in the CSF of infected IRMs were unsuccessful. However, data from other studies indicate that infection of IRMs can result in acute transmission of virus to the CNS in many animals in parallel with acute viremia (19–21). We used RT-PCR to attempt detection of virus in the CSF of the males used in the study (HP17, HP87, and JP58). In addition, some CSF samples were available from unrelated studies with ZIKV-infected male and female IRMs, and we also attempted to detect CSF-associated ZIKV in these samples. This analysis indicated that virus was present in the CNS/PNS in a majority of the animals during acute infection and was generally, but not always cleared, within two weeks (Fig. S2).

Neurological disease, including infectious neuropathy and autoimmune polyneuropathy, can result in transient or sustained elevated total protein concentration in the CSF (22). For four animals evaluated during acute infection (HP87, EL21, JI20, and JR20) with available pre-infection controls there was a significant increase in CSF protein following acute ZIKV infection as evidenced by comparison of matched preinfection controls with samples collected two weeks after infection (Fig. 4a). Pre-infection control samples were not available for the CSF samples taken from animals evaluated an extended period after infection. Two of the of the animals (HJ72 and HE27) exhibited CSF protein levels markedly higher than the samples from uninfected animals consistent with the possibility that infection can result in a longer-term increase in CSF protein in individual IRMs. However, the CSF samples from seven animals obtained from long times after infection had protein concentrations that did not differ significantly from the preinfection controls.

**Figure 4.**
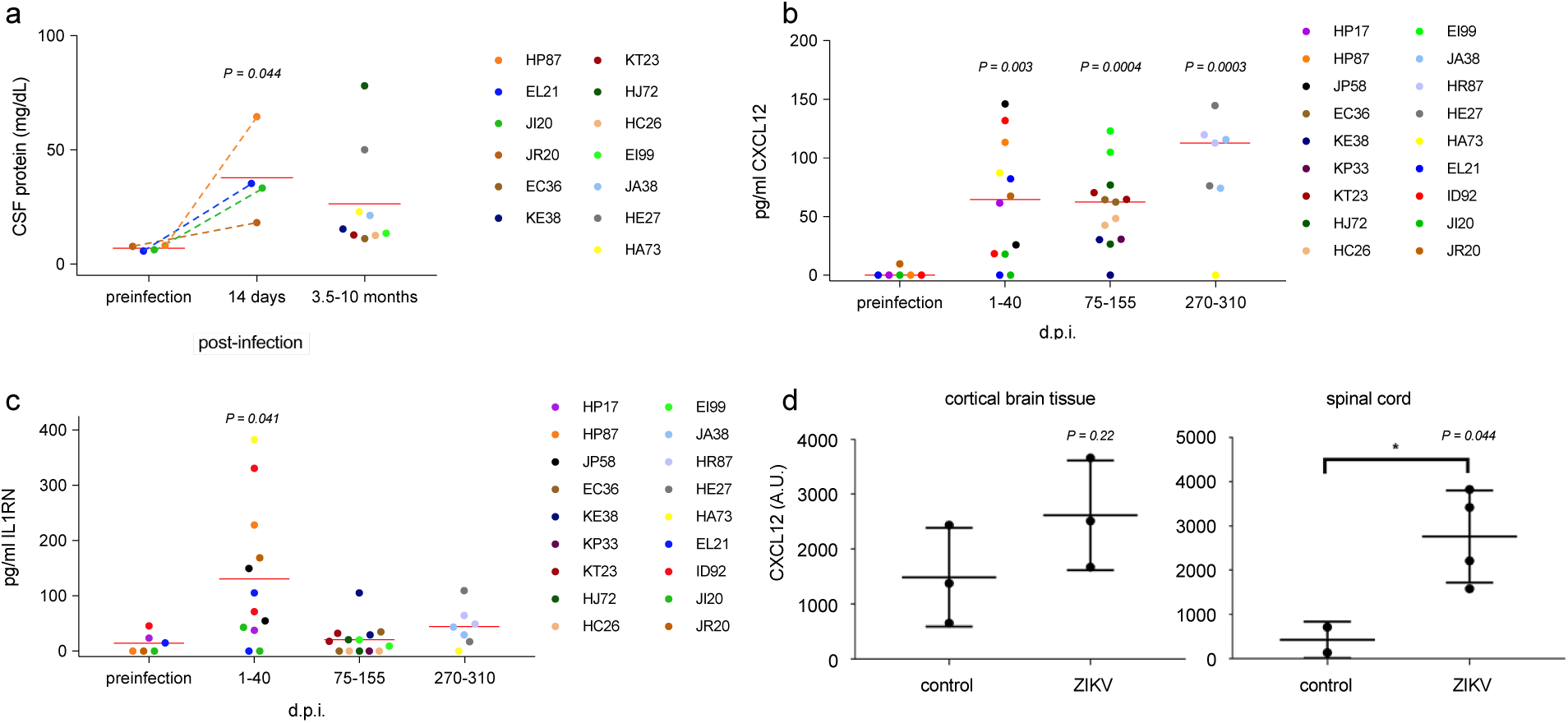
Upregulation of CXCL12 in the CSF of adult IRMs. (A) Total protein concentrations in the CSF of acutely and chronically infected IRMs. For acutely infected animals (HP87, EL21, JI20, and JR20) a two-tailed paired T-test was used to compare protein concentrations with match pre-infection samples. For samples derived from later times following infection, comparison with the pre-infection samples using a two-tailed unpaired T-test indicated that the means of the two samples were below the threshold of significance (P = 0.12). (B) and (C) Concentrations of CXCL12 and IL1RN in CSF samples following ZIKV infection, respectively. For purposes of statistical analysis samples were placed in four groups based on time of collection relative to virus infection. P values were determined by comparison to preinfection controls using two-tailed unpaired T-tests. Exact times of CSF sample collection are displayed in Fig. S4. (D) Immunohistochemistry using anti-CXCL12. CXCL12 was visualized and ImageJ was used to quantify the area of CXCL12-positive staining in 20 random images per animal. A.U. – arbitrary units.

To investigate the mechanism of ZIKV-induced neural damage and repair we carried out Luminex-based quantification of cytokine levels in the CSF in four female IRMs (EL21, JI20, JR20, and ID92) and three male IRMs (HP17, HP87, and JP58) using a macaque-specific panel designed to detect cytokines found during infection and inflammation (a list of the cytokine panel is provided in Table S1). Surprisingly, of the 37 cytokines screened and quantified in this initial analysis only two, CXCL12 and IL1RN, were significantly affected by virus infection (Fig. S3). CXCL12 is a chemokine important in multiple processes including neural repair and maintenance of BBB integrity. The IL1RN gene product is an indirect negative regulator of CXCL12; binding of virus-induced type I interferon (IFN) to its receptor, IL1R, triggers CXCL12 expression, and the IL1RN gene product binds the IL1R inhibiting IFN binding thereby blocking IFN signal transduction. We extended the analysis to all animals using CSF samples from both short and long time intervals following ZIKV infection by using a custom panel capable of quantifying CXCL12 and IL1RN. In accord with our initial experiment, CXCL12 and IL1RN were both significantly upregulated in the CSF during acute infection (Fig. 4b and c). Moreover, CXCL12 concentrations remained strikingly high in the CSF long after infection. In contrast, IL1RN markedly waned following acute infection and returned to levels similar to those observed prior to ZIKV infection.

To determine the location of CXCL12 expression in neural tissue following ZIKV infection we carried out immunohistochemistry using an anti-CXCL12 antibody. As expected, the majority of CXCL12 was detected in association with microvascular endothelia, stained with GLUT-1. Expression by vascular endothelial cells is consistent with the role of CXCL12 in maintaining BBB integrity and restriction of lymphocytes into the CNS parenchyma during homeostasis. Parallel evalutation of control and ZIKV-infected animals by semi-quantitative immunohistochemistry using an anti-CXCL12 antibody was undertaken to investigate whether the increase in CXCL12 seen in the CSF was mirrored in neural tissue. Suprisingly, we did not see a significant difference in cortical tissue between control and ZIKV-infected animals. However, expanding the analysis to spinal cord, there was a significant increase in CXCL12 staining in ZIKV infected tissue consistent with the increase in CXCL12 seen in the CSF (Fig. 4d). These results show that CXCL12 is dysregulated during Zika virus infection, and that the spinal cord is a potentially important site of action of viral infection in adults.

Positive regulators of the expression of CXCL12, as well as a spectrum of additional antimicrobial and proinflammatory cytokines, include tumor necrosis factor (TNF)(23), the proinflammatory cytokine, IL1B (24), and a soluble form of the peptide, CD40LG (25). Thus, TNF, IL1B, and CD40LG were included as targets in the custom Luminex panel we used to quantify CXCL12 and IL1RN in Fig. 4. Quantification of these potential regulators of CXCL4 indicated that all remained below the limit of detection in the CSF prior to and after ZIKV infection (Fig. S5).

## Discussion

We have shown that ZIKV causes acute and chonic neural perivascular inflammation and that ZIKV compromises BBB integrity in adult IRMs. Consequently, this is an important NHP model with high potential for elucidating facets of ZIKV-induced neuropathy in adult humans.

A striking observation is that ZIKV infection resulted in specific short- and long-term augmented expression of the chemokine CXCL12 in the CNS of adult IRMs, while other cytokines often triggered by viral infections appeared not to be expressed. Consequently, the way in which ZIKV infection induces CXCL12 is likely central to understanding ZIKV-induced neural damage and repair in adults. CXCL12 plays ubiquitous and diverse tasks in development, immunity, and repair including in the CNS. CXCL12 expression results in the recruitment or retention of CXCR4-effector cells to appropriate sites for function or homeostasis. In this regard, CXCL12 plays two roles pertinent for understanding ZIKV-associated neuropathogenesis. First, this chemokine has the capacity to regulate migration of lymphocytes through the BBB and into the CNS parenchyma (11, 12). Entry of lymphocytes across the BBB can be essential for combating viral infection in the CNS but the process can also lead to incidental neural damage through accompanying inflammation. ZIKV infection also results in augmented CXCL12 expression in monocytes(26). Trafficking of CXCL12-expressing monocytes across the BBB could further contribute to spatial skewing of the the neural CXCL12 gradient. Second, CXCL12 is important in neural repair, specifically mediating myelin restoration in the adult neural tissue through recruitment and differentiation of oligodendrocyte progenitor cells (OPCs) to effect remyelination in the CNS (7, 8), and in Schwann cell migration during repair of the PNS (9, 10). We do not yet know whether one or both of these CXCL12-dependent processes are of primary importance for understanding adult ZIKV neuropathy in the IRM.

The molecular mechanism responsible for induction of CXCL12 expression in the CNS is also unclear as we could not find overt evidence for upregulation of typical virus-induced triggers of broad cytokine induction and inflammation including INFβ, TNF, and CD40LG. The observation that CXCL12 remains upregulated in the CNS at times long after initial infection is likely significant for understanding the effect of ZIKV on the adult CNS. Sustained CXCL12 expression may be mediated by a mechanism that narrowly protracts targeted expression without concomitant expression of other cytokines. Specific post-transcription expression of CXCL12 is negatively controlled by microRNA-23a(27). It is possible that ZIKV infection results in reduced expression of this micro RNA in neural tissue. Regardless of the expression mechanism, long-term maintenance of high CXCL12 levels is likely to reflect a heretofore unrecognized chronic response of the host to ZIKV infection and may indicate the potential for virus-induced sequelae arising long after infection.

Most of the animals used in our evaluation of ZIKV on the adult CNS were females that had been infected during pregnancy. However, there does not appear to be an obvious effect of pregnancy on virus-induced CXCL12 expression in the CNS or on neuroinflammation. CSF samples taken following parturition, or from males, were not significantly different in CXCL12 levels than those obtained during pregnancy (Fig. S6). Similarly, there was not obvious correlation between time of necropsy/histological evaluation revealing inflammation and pregnancy.

We found that ZIKV infection of adult IRMs resulted in neuroinflammation with a primary outcome of high incidence meningitis. As reported here and elsewhere (19, 21), CSF-associated ZIKV appears to be cleared during acute infection. Consequently, we were surprised that neuroinflammation persisted long after infection and at times when virus is generally considered to be cleared from the CNS and other tissues. Chronic neuroinflammation could indicate that ZIKV replicates and remains in neural tissue at levels below the limit of detection. Alternatively the virus may trigger a host response marked by persistent neuroinflammation and CXCL12 expression in the absence of virus. In either case, protracted ZIKV-associated neuropathy has potentially significant clinical ramifications.

African and Asian ZIKV strains are associated with phenotypic differences in both *in vitro* replication and *in vivo* pathogenesis (28–31). The Rio-U1 ZIKV stock used in this study was a primary stock isolated from a Brazilian patient and is not adapted to cell culture. The virus forms smaller plaques on Vero cell monolayers than other strains. In addition, Rio-U1 is more pathogenic than other commonly used American strains in AG129 mice (32). It will be interesting to see whether other ZIKV strains elicit identical neuropathology to that described in this study.

While the mechanism of ZIKV-induced neuropathology in the IRM model remains to be elucidated, the neural damage and repair we observe is likely to overlap with pre-clinical or clinical ZIKV-induced GBS spectrum disease in humans. GBS and related disorders that affect the peripheral and central nervous system comprise a continuum of pathologies marked by inflammation and damage to the myelin sheath of neurons. Demyelination can arise from a constellation of genetic and environmental causes (33–38), including infection by ZIKV (39–41), and its relative Dengue virus (42, 43).

Diverse neurotropic viruses from multiple virus families gain access to the CNS parenchyma causing disruption of the BBB. Japanese Encephalitis and West Nile viruses are neurotropic viruses of the flavivirus genus, closely related to ZIKV. These viruses can replicate in the CNS, cause encephalitis, and measurably affect BBB integrity (44–49). In contrast, results from an *in vitro* BBB reconstitution model using a primary virus isolate from Thailand, and an interferon receptor-deficient murine model using Ugandan and Brazilian isolates suggests that a primary isolate of ZIKV doesn’t significantly disrupt the BBB to gain access to the CNS and that long-term damage to the BBB is minimal (50, 51). Similarly, productive ZIKV infection of human MECs in vitro does not result in cytopathic effects (44). However, our study with the NHP model indicates that ZIKV induces significant disruption of the meningeal BBB both acutely and long after infection. We do not yet know whether BBB disruption is required for CNS access in the IRM model or whether BBB damage occurs following viral CNS access and neural ZIKV replication.

## Materials and Methods

### Ethics Statement

The Indian origin rhesus macaques (IRMs)(Macaca mulatta) used in this study were housed at the TNPRC. The TNPRC is fully accredited by AAALAC International (Association for the Assessment and Accreditation of Laboratory Animal Care), Animal Welfare Assurance No. A3180-01. Animals were cared for in accordance with the NRC Guide for the Care and Use of Laboratory Animals and the Animal Welfare Act. Animal experiments were approved by the Institutional Animal Care and Use Committee (IACUC) of Tulane University (protocols P0336 and P0367). Social housing and interactive enrichment was used for all NHPs used in this study. Animal care staff conduct routine husbandry procedures (e.g., cleaning, feeding and watering), and animal care staff and veterinarians observed animals several times daily for signs of disease, pain, and distress and this information was reported to the attending veterinarian through both verbal and written communication in the animal’s health record. The Tulane University IACUC and the Division of Veterinary Medicine have established procedures to minimize pain and distress through several means. The use of preemptive and post-procedural analgesia is required for procedures that would likely cause more than momentary pain or distress in humans undergoing the same procedure. For minor procedures such as blood collection animals are anesthetized with ketamine hydrochloride (10 mg/kg IM).

### Viruses and challenge

ZIKV strain Rio U-1/2016 (16) was isolated in Rio de Janeiro, Brazil in 2016 (KU926309). Viral challenge stocks were prepared by propagating the virus in Vero cells for two passages post virus isolation(16). The stocks were quantitated by viral plaque assay. The viral stocks were diluted in Leibovitz’s L-15 and SPG media as described previously(52). All animals were challenged via the subcutaneous route, with 10^4^ PFU.

### Luminex analysis of CSF cytokines

Prior to assay, thawed serum samples were mixed well and then clarified by adding 120µl of each to Ultrafree Centrifugal Filters, pore size 0.65µm (Millipore #UFC30DV00), and centrifuged at 12,000xg for 4 minutes. Concentrations of cytokines and chemokines present in the serum were quantified using the Life Technologies Cytokine Monkey Magnetic 37-Plex Panel for Luminex™ Platform (#EPX370-40045-901, Thermo Fisher Scientific, Waltham, MA), according to manufacturer’s instructions. Assay Diluent from the kit was used to reconstitute standards and to prepare standard serial dilutions. All standards, blanks, and samples were assayed in duplicate wells. The analytes detected by this panel are indicated in Table S1. Final reactions in the microtiter plates were read on a Bio-Plex® 200 System (Bio-Rad Laboratories, Hercules, CA). Results were calculated using Bio-Plex Manager™ Software v6.1 (Bio-Rad).

### Tissue sampling and fixation

Regular peripheral blood draws and CSF collections were performed during the course of the study. At the end of the study all animals underwent a complete necropsy and tissue samples were collected in either Zinc bufferd formalin, RNA Later, RPMI media, or fresh frozen. Fixed samples were trimmed, processed, and embedding in paraffin 2 days after necropsy. Paraffin embedded tissues were cut in 5 um sections, adhered to charged glass slides, and stained routinely with hematoxylin and eosin or left unstained for immunohistochemical and immunofluorescent staining.

### Fibrinogen extravasation

The percent of vessels demonstrating fibrinogen extravasation was determined using immunofluorescence analysis with Alexa Fluor 594-labeled anti-fibrinogen and Alexa Fluor 488-labeled anti-GLUT-1 Ab, and by running linear plot profiles on the green and red channels of individual vessels and graphing the resulting numerical data in Graph Pad as dual overlay histograms. The histograms were then analyzed to determine whether the fibrinogen was above background levels outside of the two primary GLUT-1 peaks; vessels that displayed this phenotype were considered to be extravasated. A total of 25 vessels were examined from each animal via random imaging, but were required to meet the following criteria; vessels must be less than 10µm in luminal diameter and no single radius can be more than twice the length of the smallest luminal radius to ensure nearly horizontal cross sections. A Zeiss Axio Observer.Z1 fluorescence microscope was used to analyze the fluorescent labeled sections. Zeiss AxioVision Release 4.8.2 was used to capture and merge fluorescence images. Adobe Photoshop CS12.1 was also used to merge layers into a single image

### Immunohistochemistry

Immunohistochemistry was performed using anti-CXCL12. Sections were deparaffinized by incubating them for 1h at 58-60°C. After the sections were deparaffinized, they were rehydrated and pretreated for antigen retrieval by microwaving in a citrate based Antigen Unmasking Solution. Sections were washed with Tris-based saline (TBS) contain 0.05% Tween-20 for 10 min, Following, sections were treated with peroxidase blocking solution for 10 min. After washing again, sections were incubated with 5% normal goat serum in TBS for 30 minutes. Immediately following the goat serum, sections were incubated with CXCL12 antibody for 1 h at room temperature. After another wash in TBS, sections were incubated with biotinylated secondary antibody for 30 minutes. After washing the sections in TBS, sections were then incubated with Avidin Biotin peroxidase Complex for 30 min. Following another wash, sections were developed for 10 min diaminobenzidine with Mayer’s Hematoxlyin used as a nuclear counterstain. Sections were dehydrated and mounted using VectaMount. Using a Nikon Coolscope digital microscope, sections were visualized and photos were captured. Imagej was used to set a threshold level of staining and the area above this threshold was counted as positive staining with 20 random, 200x, images analyzed per animal.

### Immunofluorescence microscopy

Triple-label immunofluorescence was performed with anti-CXCL12, GLUT-1, and CD206. As described above, sections were de-paraffinized and rehydrated, followed by antigen retrieval. After washing with phosphate-buffered saline (PBS) containing 0.2% fish skin gelatin(FSG), sections were permeabilized with PBS containing 0.2% FSG and 0.1% Triton X-100 for 1 h. Following another wash, sections were incubated with 5% normal goat serum in PBS for 30 min at room temperature before incubation for 1 h at room temperature with primary antibodies diluted in PBS/FSG. After primary antibody incubation, the sections were washed in PBS/FSG and incubated with an Alexa Fluor 350-, 488-, or 594-conjugated secondary antibody in PBS/FSG for 1 h at room temperature. The sections were washed with PBS/FSG before the addition of the next primary antibody. After immunofluorescence staining, the sections were treated with 10 mM CuSO4 in 50 mM ammonium acetate buffer for 45 min to quench auto-fluorescence. The sections were rinsed in distilled water, and cover slipped with Aqua-Mount aqueous mounting medium. Confocal images were taken with a Zeiss 880 Laser scanning confocal microscope with a 100x emersion oil objective. ZenBlack and ZenBlue programs were used to capture and merge images.

### Measurement of viral RNA load (qRT-PCR)

Quantitative realtime PCR (qRT-PCR) was used for the measurement of viral loads, based on a previously validated assay(53, 54). In brief, RNA was extracted from 140 µl to 1000 µl of frozen fluids, depending on availability, using the QIAamp Viral RNA Mini Kit or QIAamp Circulating Nucleic Acid kit (Qiagen, Hilden, Germany). Total nucleic acid was eluted in two centrifugation steps with 40 µl of Buffer AVE each. A qRT-PCR reaction was then carried out with 20 µl of samples and 10 µl of primer, probes and TaqMan Fast Virus 1-Step Master Mix (Applied Biosystems, Foster City, CA). We used pre-combined probe and primers (500 nM primers and 250 nM probe; IDT Technologies, Coralville, IA). The primer and probe sequences were designed to match the sequences of the Brazilian ZIKV isolate KU321639 and were as follows: Primer 1 5’TTGAAGAGGCTGCCAGC3’; Primer 2 5’CCCACTGAACCCCATCTATTG3’; Probe 5’TGAGACCCAGTGATGGCTTGATTGC3’. The probe was double-quenched (ZEN/Iowa Black FQ) and labeled with the FAM dye (IDT Technologies, Coralville, IA). Ten-fold serial dilutions of a 401 bp *in vitro* RNA transcript encoding the ZIKV capsid gene (KU321639) starting at approximately 5 × 10^5^ RNA copies µl^−1^ were used as standards. Results were reported as the median equivalent viral RNA genomes per ml. The limit of detection was between 12 - 90 viral RNA copies ml^−1^, depending on the extracted volumes. ZIKV-positive and -negative samples were included in every run.

### Data availability

The data sets generated during and/or analysed during the current study are available from the corresponding author on reasonable request.

## Acknowledgements

This work was supported by the Bill and Melinda Gates Foundation (OPP1152818). The funders had no role in study design, data collection and analysis, decision to publish, or preparation of the manuscript.

## Supporting Information Legends

**Table S1.** Initial analysis of several animals was carried out using an NHP-specific panel designed to detect key cytokines (Cytokine/Chemokine/Growth Factor 37 Plex NHP ProcartaPlex Panel). The specific cytokines detected in the panel are listed above.

**Figure S1**. Acute viremia following ZIKV infection. (A) Viremia in eleven of the pregnant females used in the current study. (B) Viremia in the three males used in the study. Infection of nonpregnant adults results in rapid robust replication of virus as evidenced by quantification of viral RNA in serum with viral clearance from blood after about 7 to 10 days whereas viremia in pregnant females typically persists for longer times.

**Figure S2.** CSF samples collected at various times from 11 ZIKV-infected animals was analyzed using ZIKV-specific RT-PCR. Virus was detectible in the CSF in seven of the animals. Females are denoted using (x) and males by circles. None of the females used in the experiment were pregnant.

**Figure S3**. Acute infection of IRMs results in upregulation of CXCL12 (A) and IL1RN (B) in the CSF. At top, samples obtained from 14-30 days after infection are grouped to facilitate comparison with preinfection controls. P values were determined using paired two-tailed T-tests to compare CXCL12 or IL1RN concentrations following infection with that of preinfection controls. At bottom, kinetics of cytokine expression are provided with available CSF samples.

**Figure S4**. CXCL12 and IL1RN concentrations in the CSF of ZIKV-infected animals. Samples correspond to those in Fig. 4.

**Figure S5.** Quantification of TNF, IL1B, and CD40LG in the CSF of ZIKV infected IRMs. See text for description. Samples were divided into three temporal groups to facilitate comparison with Fig. 4 b and c.

**Figure S6**. This figure is reconfigured version of Fig. 4B. Points denoted with an “x” indicate that the sample was taken from a pregnant animal while points represented by a circle indicate that the sample was taken from a nonpregnant animal.

## Author Contributions

ATP, RVB, and WKK planned the studies. RVB, WKK, JBH, DGB, NJM, and BS conducted the experiments. ATP, RVB, JBH, NJM, BS, and WKK, interpreted the studies. ATP and WKK wrote the first draft. MCB provided reagents. ATP and WKK obtained funding. All authors reviewed, edited, and approved the paper.

